# Pharmacological enhancement of dopamine neurotransmission does not affect illusory pattern perception

**DOI:** 10.1101/2023.10.24.563723

**Authors:** Elke Smith, Simon Michalski, Kilian Knauth, Deniz Tuzsus, Hendrik Theis, Thilo van Eimeren, Jan Peters

**Affiliations:** Department of Psychology, Biological Psychology, University of Cologne, Cologne, Germany; Faculty of Medicine and University Hospital Cologne, Department of Nuclear Medicine, University of Cologne, Cologne, Germany; Faculty of Medicine and University Hospital Cologne, Department of Neurology, University of Cologne, Cologne, Germany

## Abstract

Psychotic symptoms and delusional belief have been linked to dopamine transmission in both healthy and clinical samples and are assumed to result at least in part from perceiving illusory patterns in noise. However, the existing literature on the role of dopamine in detecting patterns in noise is inconclusive. To address this issue, we assessed the effect of manipulating dopaminergic neurotransmission on illusory pattern perception in healthy individuals (*n* = 48, *n* = 19 female) in a double-blind placebo-controlled within-subjects design (see preregistration at https://osf.io/a4k9j/). We predicted individuals on vs. off L-DOPA to be more likely to perceive illusory patterns, specifically objects in images containing only noise. Using a signal detection model, however, we found no credible evidence that L-DOPA compared to placebo increased false alarm rates. Further, L-DOPA did not modulate measures of accuracy, discrimination sensitivity and response bias. In all cases, Bayesian statistics revealed strong evidence in favour of the null hypothesis Future studies should address possible dose-dependent effects and differential effects in healthy vs. clinical samples.

## 1 INTRODUCTION

Making sense of the world and reducing uncertainty is an essential driving force of humans trying to survive in a world full of randomness and uncertain events. By detecting patterns and meaningful relationships between stimuli or events, individuals can make predictions for the future. Recent theories conceive the brain as a system for probabilistic inference, predicting future states or events and causes of sensory input, to enable successful interaction with the environment (Friston & Stephan, 2007). Within the framework of the prediction error minimisation theory (PEM) the brain seeks to minimise the prediction errors, that is the discrepancy between predicted and actual input (Clark, 2013). The theory has been widely adopted for describing decision making, when aiming to maximise rewards (Schultz, 2016), perceptual inference (Bell et al., 2016), and social cognition, when explaining and predicting behaviour of others (de Bruin & Michael, 2021). A large body of research suggests that dopamine modulates the coding of reward prediction errors (RPEs) to enable learning, specifically phasic bursts of midbrain dopaminergic neurons code prediction errors (Schultz, 2016).

Recognising connections between stimuli or events is adaptive in many situations, as it allows making sense of the world. However, perceiving relationships between unrelated stimuli and seeing patterns in noise may be maladaptive. Seeing connections between unrelated stimuli or events prevents the formation of an accurate representation of the world. Changes in dopaminergic signalling can lead to impaired precision-weighting of prediction errors and belief updating, which then, in its most extreme form, may manifest as delusional beliefs or even psychosis (Corlett, 2018; Haarsma et al., 2021). The aberrant salience framework of psychosis describes delusional ideation as a dysfunctional computational mechanism at the neural level within a Bayesian hierarchical predictive coding framework (Kapur, 2003). The predictive coding framework describes the formation of representations to account for sensory input as a process of weighting prior beliefs against sensory inputs based its on probabilities (Sterzer et al., 2018). The psychotic state is then assumed to result from low precision of prior beliefs and dysfunctional updating of beliefs based on prior beliefs and new sensory input (Heinz et al., 2019). Within the aberrant salience framework of psychosis, dopamine is thought to modulate the salience of representations of internal states and external events. A hyperdopaminergic state would then cause aberrant assignment of salience to such representations. Delusional beliefs then reflect an individual’s cognitive effort to explain experiences of aberrant salience, while hallucinations reflect an individual’s experience of aberrantly salient internal representations (Kapur, 2003).

The link between dopamine and delusional ideations is supported by the effects of antipsychotics, which alleviate psychotic symptoms by antagonising D2 dopamine receptors (Kaar et al., 2020), and has also been confirmed using PET imaging, showing dysregulated dopamine synthesis in individuals suffering from delusions and schizophrenia (see, e.g., Cheng et al., 2020; for a review, see Rigney et al., 2021). At the conceptual level, paranormal belief, conspiratorial thinking and schizotypy are assumed to share similarities and to represent nonpathological states on a continuum converging towards delusional belief and psychosis (Denovan et al., 2018; Kreweras, 1983). These modes of cognitive processing appear to be related. For instance, conspiratorial beliefs, i.e. beliefs that certain events result from secret plots by powerful actors, were found to be positively correlated with paranormal beliefs, paranoid ideation and schizotypy (Darwin et al., 2011). Concurrently, dopamine has been linked to delusional belief and psychotic symptoms in both healthy and clinical samples, during social and perceptual decision making tasks (Barnby et al., 2020; Howes & Kapur, 2009; Krummenacher et al., 2010; Sekine et al., 2001). For instance, manipulating dopaminergic neurotransmission in controls led to changes in social attributions pertinent to paranoia, specifically D2 receptor blockade via haloperidol reduced attributions of harmful intent compared to placebo in fair and unfair social conditions, while D1 and D2 receptor stimulation with L-DOPA reduced attributions of harmful intent in fair conditions compared to placebo (Barnby et al., 2020).

For the perceptual domain, a number of studies assessed perceptual discrimination, i.e. the ability to discriminate signals and noise, in controls and individuals with hallucinations and schizophrenia (Bentall & Slade, 1985; Ishigaki & Tanno, 1999; Krummenacher et al., 2010). Specifically modelling response bias and discrimination sensitivity (d-prime), the studies yielded mixed results, however. Early studies report a more liberal criterion, i.e. tendency to identify signals, but no difference in discrimination sensitivity, for individuals with high compared to low predisposition to hallucination in auditory signal detection (Bentall & Slade, 1985), and decreased discrimination sensitivity of patients with schizophrenia with concomitant auditory hallucinations compared to controls in a visual continuous performance test (Ishigaki & Tanno, 1999). In contrast, a more recent study reports individuals with paranormal beliefs to exhibit a response strategy of favouring false alarms over misses, and individuals sceptical about paranormal phenomena showing the reverse strategy. Enhancing dopaminergic neurotransmission lowered discrimination sensitivity compared to placebo in the group of sceptics, but not believers (Krummenacher et al., 2010).

Dopamine has been assumed to modulate neuronal activity in such a way that signal transmission is less distorted by noise (Vander Weele et al., 2018; Walter & Spitzer, 2003). However, considering the above findings and psychosis as a hyperdopaminergic state characterised by poor ability to discriminate relevant and irrelevant, and internal and external stimuli (Chu et al., 2021; Morris et al., 2013), this assumption falls short and the findings on the exact relationship between dopamine and perceptual discrimination remain inconclusive. The inconsistencies might be related to differences in sample characteristics, for instance to differences in schizophrenia symptomatology, or related to the domain under study, such as auditory or visual. Further, most of the studies published rely on rather small subgroup sample sizes (Barnby et al., 2020; Ishigaki & Tanno, 1999; Krummenacher et al., 2010). To contribute to resolving these inconsistencies, we studied the influence of enhancing dopaminergic neurotransmission with the dopamine precursor L-DOPA on illusory pattern perception in healthy individuals, modelling discrimination performance for detecting objects in images with signal detection theory. We predicted participants on vs. off L-DOPA to exhibit increases in false alarms, i.e. to perceive more illusory patterns, specifically objects in images containing only noise (see preregistration at https://osf.io/a4k9j/). With regard to response bias, discrimination sensitivity (d-prime) and accuracy, our hypotheses were non-directional.

## 2 METHODS

### 2.1 Sample

A subset of *n* = 49 out of *N* = 76 participants from a larger pharmacological study performed the experiment. One participant was excluded due to side effects (nausea and vomiting) and did not complete the task. The final sample included 48 participants, including 19 women, all right-handed, aged 25 to 40 (*M* = 28.27). The participants were recruited through university bulletins, mailing lists and by word-of-mouth recommendation. For practical reasons, only a subset of participants completed the SPT. Therefore, no task-specific a priori power calculation was carried out. A post-hoc power analysis (paired sample test with *G*Power*, version 3.1.9.7, Faul et al., 2009) yielded a power of 0.39 to detect a small effect (*d* = 0.2), power of 0.96 to detect a medium effect (*d* = 0.5), and a power of > 0.99 to detect a large effect (*d* = 0.8). All participants had normal or corrected-to-normal vision, German as first language (or profound German language skills), and all women were taking hormonal contraceptives. General exclusion criteria for study participation were strongly impaired vision or strabismus, participation in other studies involving medications, intake of non-prescription and prescription drugs, pregnancy, acute infections, alcohol or drug intoxication or abuse, psychiatric disorders (past or current), neurological disorders, metabolic disorders, internal diseases, chronic pain syndrome, complications of anaesthesia, and strong emotional burden or physical stress during the study period. Exclusion criteria considering contraindications regarding the intake of L-DOPA were hypersensitivity to L-DOPA or benserazide, intake of non-selective monoamine oxidase inhibitors, metoclopramide or antihypertensive medication (e.g., reserpine), disorders of the central dopaminergic system, e.g., Lewy body dementia or Parkinson’s disease, increased intraocular pressure (glaucoma), and breast feeding.

### 2.2 Procedure

After passing a medical examination by a physician to check for contraindications, the participants were invited to three testing sessions. During the first session, the participants underwent a baseline screening for putative dopamine proxies, specifically working memory capacity, spontaneous eye blink rate, and impulsivity. After the baseline screening, the participants completed two identical experimental sessions on separate days with an interval of about a week between the sessions. 30 minutes prior to testing, the participants received a non-distinguishable tablet containing either 150 mg L-DOPA, a dopamine precursor, and benseracide, a peripheral decarboxylase inhibitor, or placebo medication and then completed an intertemporal choice task and a reinforcement learning task (see preregistration at https://osf.io/a4k9j/). A subset of participants additionally completed a perceptual discrimination task (see section 2.3) approximately 75 minutes after intake of the tablet. The data from the baseline screening, intertemporal choice task and reinforcement learning task will be reported elsewhere. The study was realised as double-blind placebo-controlled within-subjects design. Polling the participants showed that they could not guess the correct order of the experimental sessions χ^2^ (1, *N* = 48) = 0, *p* = 1.0.

### 2.3 Snowy Pictures Task

We used a modified pen-and-paper version of the snowy pictures task (Ekstrom & Harman, 1976; Whitson & Galinsky, 2008). The task contains 24 grainy images. Some of the images contain hard-to-detect embedded objects (for instance a chair and a knife), some contain only noise (12 images with objects, 12 images with noise). The participants were asked to denote whether or not an object is present in the image, and if so, what object it is. They were instructed to complete the task as fast as possible without sacrificing accuracy. The participants completed two different versions of the task under placebo and L-DOPA, respectively, in counterbalanced order (12 images per session).

### 2.4 Data analysis

To assess the influence of enhancing dopaminergic transmission on the detection of objects in images, we calculated accuracies, false alarm rates, and the signal detection theory measures d-prime and response bias per condition (placebo and L-DOPA). D-prime is an index for the ability to disentangle signal from noise, with higher values reflecting greater discriminability, while the response bias reflects the tendency towards responding ‘yes’ or ‘no’. Negative values indicate a liberal response criterion (response bias towards responding ‘yes’), while positive values indicate a conservative response criterion (response bias towards responding ‘no’). D-prime and response bias were computed participant- and condition-wise in *MATLAB* (version R2023a) based on Stanislaw & Todorov (1999). Significant Shapiro-Wilk tests for accuracy *W* = .928, *p* = .006, false alarm rate *W* = .909, *p* = .001 and response bias differences *W* = .934, *p* < .009 indicated that differences between the matched pairs were not normally distributed. Therefore, we used non-parametric Bayesian Wilcoxon signed-rank tests as implemented in *JASP* (version 0.17.2.1) to compare the group means of accuracy, false alarm rate and response bias between conditions. Since D-prime was normally distributed (*W* = .971, *p* = .270), we used a Bayesian paired samples t-Test to compare d-prime between conditions. The posterior distributions were obtained using Markov-Chain Monte-Carlo (MCMC) sampling with 5 chains and 1000 samples, using a Cauchy distribution with scale = 2 as prior. We report Bayes factors to evaluate evidence in favour of the null hypothesis (non-directional: *BF*_*01*_, directional: *BF*_*0-*_).

## 3 RESULTS

On average, and across both conditions (placebo and L-DOPA), participants correctly detected objects or correctly rejected noise in 81.77% of all images. The average response bias across both conditions of 0.65 indicates an overall tendency towards responding ‘no’ (i.e. no object identified in an image). Under L-DOPA, participants made false alarms in 11.28% of all images, i.e. identified objects in images containing only noise, compared to 12.15% in the placebo condition (see Table 1 and Figure 1 for accuracy, hits, false alarms, d-prime and response bias per drug condition). Testing for group differences in accuracy, false alarm rate, response bias and d-prime between the conditions, revealed that the null effect of no difference in accuracy under L-DOPA and placebo was 14 times more likely than a difference between the conditions (*BF*_*01*_ = 14.48). Likewise, a null effect for the false alarm rate between L-DOPA and placebo was 20 times more likely than an increased false alarm rate under L-DOPA vs. placebo (*BF*_*0-*_ = 20.25). Furthermore, Bayesian analyses revealed evidence in favour of the null hypothesis for the response bias (*BF*_*01*_ = 6.37), and d-prime (*BF*_*01*_ = 11.46).

**Table 1:**
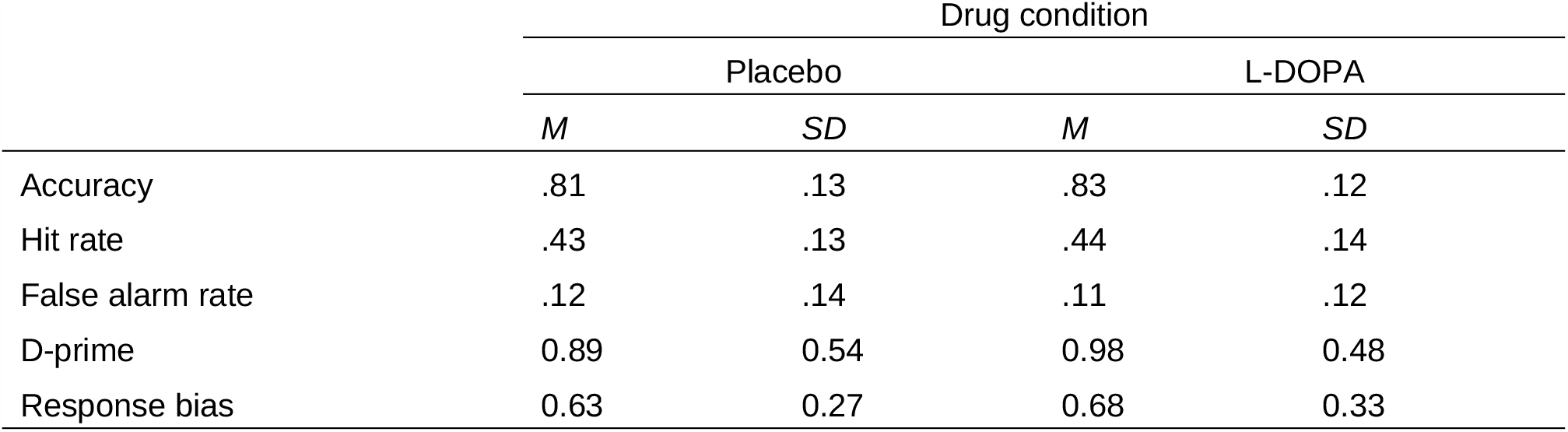
Means and standard deviations for task performance per drug condition.

**Figure 1.**
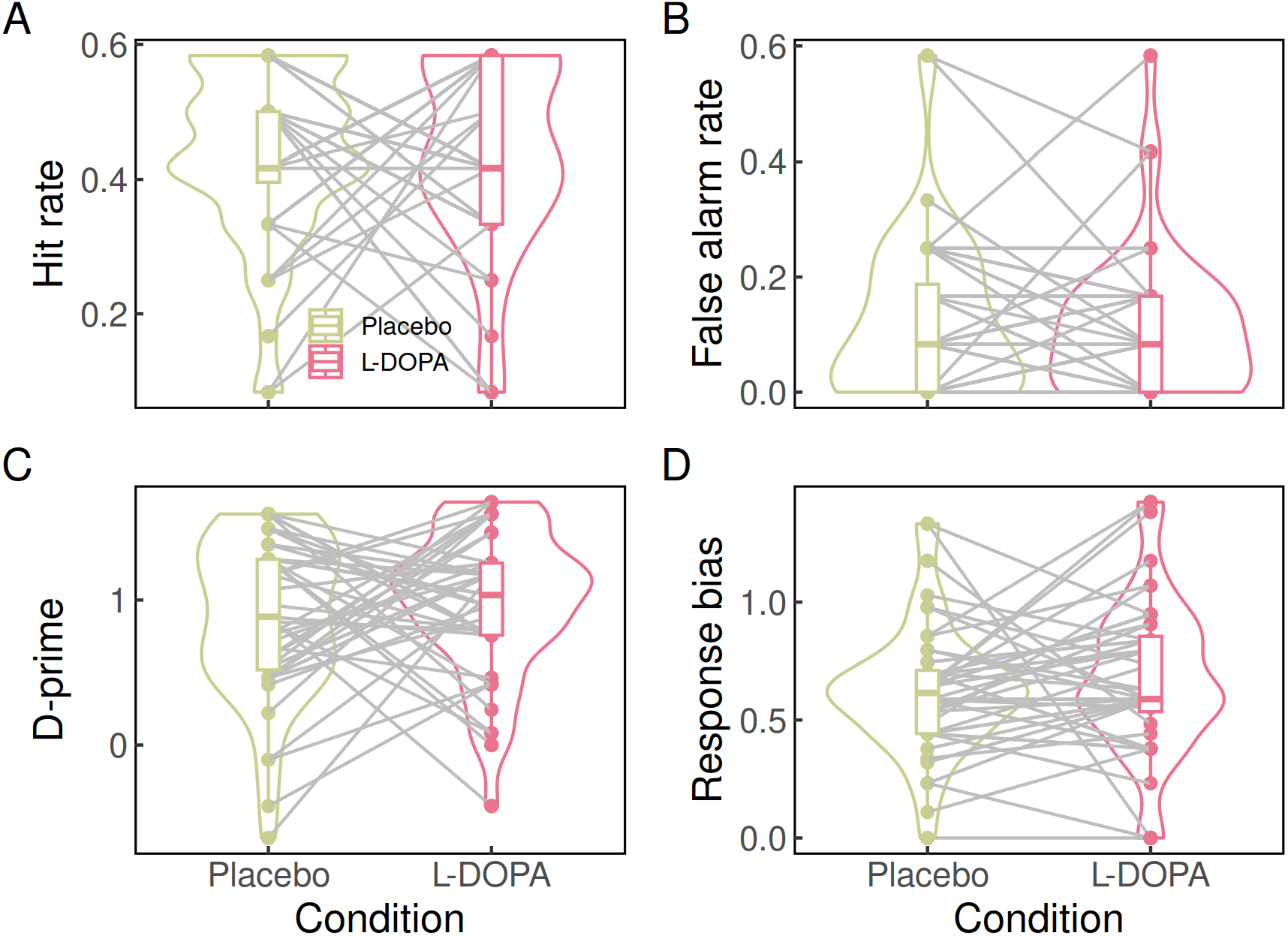
Distribution of accuracy (A), false alarm rate (B), d-prime (C) and response bias (D) per condition. Horizontal line: median, box: first and third quartiles, lower whisker: lowest value not less than 1.5*IQR (inter-quartile range) from the first quartile, upper whisker: highest value no further than 1.5*IQR from the third quartile, dots: outliers (data points outside the lower and upper whisker). For the hit rate and false alarm rate (panel A and B, respectively), there are overlapping data points.

## 4 DISCUSSION

We studied the effects of enhancing dopamine neurotransmission using the dopamine precursor L-DOPA on illusory pattern perception using a visual object identification task. Participants completed two versions of the modified Snowy Pictures Task (Ekstrom & Harman, 1976; Whitson & Galinsky, 2008) under placebo and L-DOPA, respectively. Applying signal detection theory, our hypothesis that participants on L-DOPA would be more likely to perceive illusory patterns, specifically objects in images containing only noise, was not confirmed. In contrast, Bayesian analyses revealed strong evidence in favour of the null hypothesis for false alarm rates. Likewise, for accuracy, discrimination sensitivity and response bias, Bayesian analyses revealed evidence in favour of the null hypothesis.

Changes in discrimination sensitivity and social attributions relevant to paranoia following dopaminergic modulation have previously been reported in perceptual and social decision making in healthy samples (Barnby et al., 2020; Krummenacher et al., 2010). The involvement of dopamine in delusional ideation is further substantiated by the efficacy of antipsychotic treatment (Kaar et al., 2020), and a study assessing perceptual discrimination in controls and individuals with hallucinations and schizophrenia reported lower discrimination sensitivity in patients with auditory hallucinations (Ishigaki & Tanno, 1999). The present null effect of L-DOPA on illusory pattern perception in the current study may be for several reasons. First, earlier studies reporting a relationship between dopamine and discrimination sensitivity or paranoid inferences rely on rather low sample sizes (30 participants in within-subjects design in Barnby et al., 2020, 20 participants per belief group in between-subject design in Krummenacher et al., 2010, 11 participants per patient group in Ishigaki & Tanno, 1999). Low sample sizes increase the variance in effect sizes even under the null, such that previously reported findings may have been false positives. Alternatively, the present null effect may be related to dose-dependent effects, or a single dose may have not been sufficient to elicit detectable changes in pattern perception. The effect of pharmacological DA manipulation might also depend on interindividual differences, such as a predisposition to delusional thinking, belief in the paranormal (Krummenacher et al., 2010), magical ideation (Mohr et al., 2006), and predisposition to hallucinations (Bentall & Slade, 1985). For instance, L-DOPA increased semantic priming only in participants with high magical ideation (due to longer response times for unrelated prime-target pairs), and participants with high magical ideation under placebo performed comparable to participants with low magical ideation under L-DOPA (Mohr et al., 2006). It is further conceivable that dopamine is related to delusional beliefs, while manifesting itself only in the pathological state or in individuals scoring high on schizotypy (Mohr & Ettinger, 2014). Lastly, the nature of the task may not suitable to detect dopamine-related changes in illusory pattern perception, since it covers the visual domain only, and requires no inferences about events or social intentions.

### Limitations

We did not assess the participants’ baseline dopamine synthesis capacity, and their predisposition to delusional thinking or paranormal belief. The participants in our sample may not have had such a predisposition and a single dose of L-DOPA may therefore not have been sufficient to induce such effects. Also, since overall accuracy was rather high, and false alarm rates rather low, the task may be susceptible to ceiling effects.

### Conclusion and perspectives

We assessed the effect of enhancing dopaminergic neurotransmission on illusory pattern perception. There was no evidence that L-DOPA, compared to placebo, increases the detection of patterns is noise. Rather, Bayesian analyses provided strong evidence in favour of the null hypothesis. Future studies should control for predisposition to delusional thinking, belief in the paranormal or magical ideation and may assess changes in pattern perception across different domains (e.g., auditory and visual).

## Author contributions

Conceptualisation: J. P., E. S.

Data collection: E. S., K. K., D. T., H. T., T. E.

Methodology: E. S., S. M.

Formal analysis: E. S., S. M.

Writing - Original Draft: E. S., S. M.

Writing - Review & Editing: E. S., S. M., K. K., D. T., H. T., T. E., J. P.

Project administration: E.S., J. P.

Supervision: J. P.

## Acknowledgements

Lea Kemalides, Hannah Hacker, Emily Burlon

## Funding

This work was funded by the Deutsche Forschungsgemeinschaft (DFG, project no. PE1627/5-1). H.T. was supported by the Cologne Clinician Scientist Program (CCSP) of the Faculty of Medicine of the University of Cologne, funded by the Deutsche Forschungsgemeinschaft (DFG, German Research Foundation, project no. 413543196).

## Ethics statement

The study was approved by the local ethics committee of the Faculty of Medicine of the University of Cologne, Germany.

## Data availability statement

The data that support the findings of this study are openly available an https://osf.io/m5u6v/.

## Notes

https://osf.io/a4k9j/

### Competing Interest Statement

The authors have declared no competing interest.

### Summary of Updates

Added information on excluded participants.

